# Identification of HIV-Transmitting Sub-Epithelial Mononuclear Phagocytes in Human Anogenital and Colorectal Tissues

**DOI:** 10.1101/2020.05.26.117408

**Authors:** Jake W Rhodes, Rachel A Botting, Kirstie M Bertram, Hafsa Rana, Heeva Baharlou, Erica E Longmuir-Vine, Peter Vegh, James Fletcher, Thomas R O’Neil, Grant P Parnell, J Dinny Graham, Najla Nasr, Jake J K Lim, Laith Barnouti, Peter Haertsch, Martijn P Gosselink, Angelina Di Re, Grahame Ctercteko, Gregory J Jenkins, Andrew J Brooks, Ellis Patrick, Scott N Byrne, Muzlifah A Haniffa, Anthony L Cunningham, Andrew N Harman

## Abstract

Tissue mononuclear phagocytes (MNP) are specialised in pathogen detection and antigen presentation. They are the first cells of the immune system to encounter HIV and play a key role in transmission as they deliver the virus to CD4 T cells, which are the primary HIV target cell in which the virus undergoes replication. Most studies have investigated the role that epithelial MNPs play in HIV transmission but, as mucosal trauma and inflammation are strongly associated with HIV transmission, it is also important to examine the role that sub-epithelial MNPs play. Sub-epithelial MNPs are present in a diverse array of subsets which differ in their function and the pathogens they detect. Understanding how specific subsets interact with HIV and deliver the virus to CD4 T cells is therefore of key importance to vaccine and microbicide development. In this study we have shown that, after topical application, HIV can penetrate to interact with sub-epithelial resident myeloid cells in anogenital explants and defined the full array of MNP subsets that are present in all the human anogenital and colorectal sub-epithelial tissues that HIV may encounter during sexual transmission. In doing so we have identified two subsets that preferentially take up HIV, become infected and transmit the virus to CD4 T cells; CD14^+^CD1c^+^CD11c^+^ monocyte-derived dendritic cells and langerin-expressing dendritic cells 2 (DC2).

## Introduction

There is still no cure or vaccine for HIV / AIDS and 37 million people remain infected. Antiretroviral therapy (ART) is efficient at controlling infection but is a lifelong treatment which is costly to manage and associated with toxicities. Only 57% of HIV^+^ individuals receive ART and there are 1.8 million new infections each year. ART can be given to healthy ‘at risk’ individuals as pre-exposure prophylaxis (PrEP) which has shown to be effective in reducing transmission. However, this is not a universal solution because of poor access to PrEP in low income countries and variable uptake in Western countries. Furthermore, the effects of long-term administration of PrEP to healthy individuals are unknown and can be associated with decreased condom use^1^, increased sexually transmitted infections and concomitant genital tract inflammation^2^, enhancing HIV transmission^3-6^, especially in sub-Saharan Africa. Furthermore, PrEP regimens have recently been shown to be ineffective in the context of an inflamed mucosa^7-9^. Therefore, an effective vaccine and cure are still needed.

HIV is now transmitted sexually in almost all cases. In order to develop a vaccine (or more effective PrEP regimens) the precise definition of the initial HIV target cells in the anogenital mucosa is necessary, especially mononuclear phagocytes (MNP). These consist of Langerhans cells (LC), dendritic cells (DC) and macrophages which express the HIV entry receptors CD4 and CCR5 allowing them to be directly infected. Importantly, they also express a large repertoire of lectin receptors including C-Type Lectins (CLR) and Sialic acid-binding immunoglobulin-type lectins (Siglec), many of which can bind HIV and mediate endocytic uptake of the virus^10-15^. As professional antigen presenting cells, LCs and DCs play a critical role in HIV transmission by transferring the virus to CD4 T cells^13,16,17^ where it replicates resulting in CD4 T cell death and depletion and consequent immunosuppression. Epidermal Langerhans cells (LC) have been shown to take up HIV and transfer it to T cells in vagina, cervix and foreskin^18-20^ and we and others have recently shown that epidermal DC populations also participate in this process^21,22^. We also previously showed that HIV transfer to CD4 T cells occurs in two successive phases from epidermal LCs^13^, epidermal CD11c^+^ DCs^21^ and *in vitro* derived monocyte-derived (MDDC)^23,24^. The first phase of transfer occurs within 2 hours and is dependent on lectin mediated uptake of the virus and declines rapidly with time. The second phase occurs from 72 hours onwards and increases with time as newly formed virions bud off from the surface of cells that have become productively infected via CD4/CCR5 mediated entry into viral synapses^17^.

Mucosal trauma which breaches anogenital and colorectal epithelium and associated inflammation are likely to enhance HIV acquisition as it also allows the virus direct access to deeper target cells in the underlying lamina propria. Despite this, the role of lamina propria MNP subsets in HIV transmission has been largely understudied. These MNPs are present in several distinct subsets which include cDC1^25,26^ and cDC2^27^. In addition, there are several subsets of CD14^+^ cells^28-32^ which include autofluorescent tissue resident macrophages^27^ and non-autofluorescent CD14^+^ cells which have been conventionally referred to as DCs^27^ which have an established role in transmitting HIV to CD4 T cells in both intestinal^33-35^ and cervical^15^ tissue. However in healthy tissue CD14^+^CD1c^-^ cells have recently been redefined as monocyte-derived macrophages (MDM)^36^. This makes their role in HIV transmission puzzling as macrophages are very weak antigen presenting cells for naïve T cells and do not migrate to lymph nodes and thus are less likely to deliver the virus to CD4T cells than DCs. In inflamed tissue CD14^+^CD1c^+^ cells have been described which transcriptionally align with DCs^37,38^ and have been defined as *in vivo* MDDCs^32^. Recently similar cells have also been described in healthy lung tissue^39^.

In this study we have thoroughly defined the MNP subsets that are present in the sub-epithelium (lamina propria and dermis) of all human anogenital and colorectal tissues and found two cell subsets that are key players in HIV uptake and transmission; CD14^+^CD1c^+^ MDDCs and langerin^+^ cDC2. Compared to other MNPs both subsets took up HIV more efficiently and became more infected. We found that *ex vivo* CD14^+^CD1c^+^ MDDCs were most efficient at transferring HIV to CD4 T cells at late time points which correlated with their high CCR5 expression and that langerin^+^ cDC2 transferred the virus most efficiently at early time points.

## Materials and Methods

### Sources of Tissues and Ethical Approval

This study was approved by the Western Sydney Local Area Health District (WSLHD) Human Research Ethics Committee (HREC); reference number HREC/2013/8/4.4(3777) AU RED HREC/13/WMEAD/232. Healthy human tissue was obtained from a range of plastic surgeons and written consent was obtained from all donors.

### Tissue Processing

MNP were isolated from abdominal tissue using our optimised collagenase-based digestion process^21,40^. Skin was collected immediately after surgery, stretched out and sectioned using a skin graft knife (Swann-Morton,Sheffield, United Kingdom) and the resulting skin grafts passed through a skin graft mesher (Zimmer Bionet, Warsaw, IN, USA). The meshed skin was placed in RPMI1640 (Lonza, Switzerland) with 0.14U/ ml dispase (neutral protease, Worthington Industries, Columbus, OH, USA) and 50ug/mL Gentamicin (Sigma-Aldrich, St Louis, MO, USA) and rotated at 4°C overnight. The skin was then washed in PBS and dermis and epidermis were mechanically separated using fine forceps. Dermal tissue was cut into 1-2 mm pieces using a scalpel. Dermal and epidermal tissue was then incubated separately in RPMI1640 containing 100U/ml DNase I (Worthington Industries) and 200U/ml collagenase Type IV (Worthington) at 37°C for 120 minutes in a rotator. The cells were then separated from undigested dermal and epidermal tissue using a tea strainer. The supernatants were then passed through a 100μm cell strainer (Greiner Bio-One, Monroe, NC, USA) and pelleted. The cell pellet was then passed again through a 100μm cell strainer, and incubated in MACS wash (PBS (Lonza) with 1% (v/v) Human AB serum (Invitrogen, Carlsbad, CA, USA) and 2 mM EDTA (Sigma-Aldrich) supplemented with 50U/mL DNase for 15 minutes at 37°C. The epidermal suspension was spun on a Ficoll-Paque PLUS (GE Healthcare Life Sciences, Little Chalfont, United Kingdom) gradient and the immune cells harvested from the Ficoll-PBS interface. Dermal cells were enriched for CD45-expressing cells using CD45 magnetic bead separation (Miltenyi Biotec,San Diego, CA, USA). Cell suspensions were then counted and/or labelled for flow cytometric phenotyping of surface expression markers or for flow sorting. This protocol was modified for anogenital tissues as follows: for Skin and Type II mucosa (labia, foreskin, glans penis, fossa navicularis, vagina, ectocervix and anal canal), small shallow scalpel cuts were made to the epithelial surface before overnight Dispase II treatment; for Type I mucosa (endocervix, penile urethra, rectum and colon) no Dispase treatment was required and tissue was digested using two successive 30 minute digestions with Collagenase Type IV. For spontanteous migration assays, following mechanical separation of dermis and epidermis tissue was cultured for 48hrs in RPMI 10% FCS, 50 U/ml DNase I and 25µg/ml gentamicin. Isolated cells were then treated as above.

### Preparation of *in vitro* Monocyte Derived Dendritic Cells and Macrophages

CD14^+^ monocytes were derived from human blood and cultured for six days with interleukin (IL)-4 and granulocyte-macrophage colony-stimulating factor (GM-CSF) to produce *in vitro* derived MDDC or in human serum to generate *in vitro* derived MDM as described previously^23,41-44^.

### Flow Cytometry and Sorting

Cells were labelled in aliquots of 1×10^6^ cells per 100μl of buffer, according to standard protocols. Nonviable cells were excluded by staining with Live/Dead Near-IR dead cell stain kit (Life Technologies, Carlsbad, CA, USA). Flow cytometry was performed on Becton Dickenson (BD, Franklin Lakes, NJ, USA) LSRFortessa flow cytometer and data analysed by FlowJo (Treestar). Fluorescence Activated Cell Sorting (FACS) was performed on a BD FACS Influx (100μm nozzle and 20 pounds/square inch). Sorted cells were collected into FACS tubes containing RPMI culture media supplemented with 10µM HEPES (Gibco, Waltham, MA, USA), non-essential amino acids (Gibco), 1mM sodium pyruvate (Gibco), 50µM 2-mercaptoethanol (Gibco), 10µg/ml gentamycin (Gibco) and 10% (v/v) FCS (from herein referred to as DC culture media. The antibodies were purchased from BD, Miltenyi, Bio Legend (San Diego, CA, USA), Beckman Coulter (Brea, CA, USA), eBioscience (San Diego, CA, USA) and R&D Systems (Minneapolis, MN, USA) as follows; BD: CD45 BV786 (HI30), HLA-DR, BUV395 (G46-6), CD1a BV510 (HI149), CD14 BUV737 (M5E2), CD141 BV711 (1A4), CD33 (Siglec-3) APC (WM53), CD161 (Clec5B) APC (DX12), CD184 (CXCR4) PE (12G5), CD103 PE (Ber-ACT8), CD4 APC (RPA-T4), CD195 (CCR5) PE (2D7), CD209 (DC-SIGN) APC (DCN46), CD80 PE (L307.4), CD83 APC (HB15e), CD86 APC (2331 (FUN-1), CD371 (Clec12A) AF647 (50C1), Mouse IgG1 APC, Mouse IgG2b APC, Mouse IgG1 PE, Mouse IgG2a PE. Miltenyi: CD14 Vioblue (TUK4), Clec7A (Dectin-1; CD369) PE (REA515), Clec9A (CD370) PE (8F9), CD1c (BDCA1) PE-Vio770 (AD5-8E7), CD141 APC (AD4-14H12), CD207 (Langerin; Clec4K) Vioblue (MB22-9F5), Mouse IgG2a PE, Mouse IgG1 APC. Bio Legend: CD169 (Siglec-1) PE (7-239), DEC205 PE (HD30), XCR1 APC (S15046E). Beckman Coulter: CD207 PE (DCGM4), CD206 (MR) PE (3.29B1.10), Mouse IgG1 PE (679.1Mc7). eBioscience: CD91 eFluor660 (A2MR-a2), CD172a (SIRPα) PerCP-eFluor710 (15-414), Mouse IgG1 eFluor660. R&D Systems: CD299 (L-SIGN; DC-SIGNR) PE (120604), CD367 (Clec4A; DCIR) PE (216110), CD368 (Clec4D) PE (413512), Clec5A APC (283834), Clec5C APC (239127), Clec6A (Dectin-2) APC (545943), CD301 (Clec10A) PE (744812), Clec14A APC (743940), Siglec-9 APC (191240), Siglec-16 APC (706022), Mouse IgG1 PE, Mouse IgG2b PE, Mouse IgG2b APC, Mouse IgG2a APC. For HIV detection cells were incubated with Cytofix/Cytoperm solution (BD) for 15 minutes at room temp and washed with permeability wash (1% BSA, 0.05% Saponin, 0.1% NaN_3_ in PBS). Samples were then blocked with 10% HuAb serum for 10 minutes and stained with two antibody clones to HIV p24, KC57 PE (Beckman Coulter) and 28B7 APC (Medimabs, Canada).

### RNA seq

Total RNA (1ng in 2μl) was reverse transcribed and amplified using the Smart-seq2 protocol of Picelli et al^45,46^. Transcripts were first mixed with 1μl of 10μM anchored oligo-dT primer (5’-AAGCAGTGGTATCAACGCAGAGTACT_30_VN-3’, and 1μl dNTP mix (10mM) and heated to 72°C for 3 minutes then placed on ice immediately. A first strand reaction mix containing Superscript II, TSO primer (5’-AAGCAGTGGTATCAACGCAGAGTACATrGrG+G-3’, 1uM, 1M betaine, 6mM MgCl_2_ 5mM DTT, RNaseOUT (10U) and 2ul 5X first strand buffer, was added to each sample. Transcripts were reverse transcribed by incubating at 42°C for 60 minutes, followed by 10 cycles of 50°C, 2 minutes and 42°C, 2 minutes. The resulting cDNAs were subsequently amplified by adding 12.5μl KAPA HiFi 2X HotStart ReadyMix, 10μM ISPCR primer (0.25μl, 5’-AAGCAGTGGTATCAACGCAGAGT-3’) and 2.25μl H_2_O and heating to 98°C, 3 minutes, followed by 15 cycles of 98°C, 20 seconds, 67°C, 15 seconds, 72°C, 6 minutes. After a final incubation at 72°C, 5 minutes, samples were held at 4°C until purification. Samples were purified by addition of Agencourt AMPure XP beads (1:1 v/v), 8 minutes, room temperature, followed by magnetic bead capture and two washes with 200μl 80% ethanol. Beads were dried and cDNAs were eluted into 17.5μl elution buffer EB. Products were checked on Agilent Bioanalyser High Sensitivity DNA chips and yield was estimated by Quant-iT PicoGreen dsDNA Assay. Sequencing libraries were prepared from 2ng product using Nextera XT reagents in accordance with the supplier’s protocol except that PCR was limited to 8 cycles to minimize read duplicates, and final library elution was performed using 17.5μl elution buffer EB. Libraries were pooled and sequenced on the Illumina HiSeq 2500 platform generating 50 base pair single-end sequences. Raw sequence data was aligned to the UCSC human reference genome (hg19) using the TopHat2 software package^47^. Aligned sequencing reads were summarised to counts per gene using the Read Assignment via Expectation Maximisation (RAEM) procedure^48^ and reads per kilobase per million mapped reads (RPKM) values were calculated in SAMMate (v 2.6.1)^48^.

### HIV Transfer Assay

*Ex vivo*-derived dermal mononuclear phagocyte (MNP) subsets were tested for their ability to transfer HIV to activated T cells as described previously for Epidermal MNPs^13, 21^. Cells were liberated from abdominal skin as described above and individual subsets were FACS isolated. A minimum of 1×10^4^ and 3×10^4^ cells were cultured with HIV_Bal_ for 1^st^ Phase and 2^nd^ phase transfer assays respectively, at an MOI of 1 for 2 hours before being washed off three times with PBS. For 1^st^ Phase transfer assays, JLTR cells (CD4 T cells which express GFP under the HIV promoter) were co-cultured at a ratio of 4:1 (T cells:MNPs) for 96 hours in DC culture media. For 2^nd^ phase transfer assay, cells were cultured for 96 hours before addition of JLTR cells. Flow cytometry was used to determine the percent of GFP^+^ JLTRs.

### Secreted HIV Infectivity Assay

Cell supernatants from 2^nd^ phase transfer assays were taken prior to the addition of JLTRs (96 hours after virus had been washed off). Supernatants were cultured with TZMBL cells (HeLa cell derivatives expressing high levels of CD4 and CCR5 and containing the β-galactosidase reporter gene under the HIV promoter) for a further 72 hours on 96-well flat-bottom plates. Supernatants were removed and TZMBL development solution added (0.5M potassium ferricyanide, 0.5M potassium cyanide, 50mg/ml X-gal) and incubated for 1 hour to detect LTR β-galactosidase reporter gene expression. Development solution was removed, and cells fixed with 4% (v/v) PFA (diluted in PBS) for 15 minutes. Infectivity was quantified using an EliSpot plate reader (AID, Strasberg, Germany).

### HIV Uptake Assay

*Ex vivo* dermal MNP subsets were tested for their ability to take up HIV virions. Cells were liberated from abdominal tissue through enzymatic digestion and CD45^+^HLA-DR^+^ cells were FACS isolated. Sorted cells were cultured for 2 hours with HIV_Bal_ or HIV_Z3678M_ at an MOI of 3 before being washed of 3x with PBS. Cells were labelled for flow cytometric analysis as described above and analysed to determine the percent of dual p24^+^ cells. For Siglec-1 blocking assays, cells were pre-treated with 20µg/ml Siglec-1 mAb (Invitrogen, HSn 7D2) for 30 minutes before HIV was added.

### Inner Foreskin HIV Explants

Inner foreskin explants were infected with HIV_Bal_ as previously described^21^. Briefly, small scalpel cuts were punctured into the mucosal surface of the tissue before cloning cylinders (8×8mm, Sigma-Aldrich) were adhered to the tissue using surgical glue (B Braun, Germany). Explants were cut around the cloning cylinder and placed onto gel foam sponge (Pfizer, New York, NY, USA) soaking in DC culture media in a 24-well plate. HIV was diluted in 100µl of PBS at a TCID_50_ of 3500 and then added to cloning cylinders. 100µl of PBS was added to mock samples to prevent tissue drying out. Explants were cultured for 2-3 hours before virus/PBS was removed and cylinders washed out 3x with PBS. After removal of cloning cylinders, tissue explants were placed in 4% PFA (Electron Microscopy Sciences, Hatfield, PA, USA) for 24 hours before paraffin embedding.

### RNAscope and immunofluorescent staining of tissue

All microscopy staining was carried out on 4µm paraffin sections. Sections were baked at 60°C for 45 minutes, dewaxed in xylene followed by 100% ethanol and dried. Detection of HIV RNA was achieved using the RNAscope 2.5HD Reagent Kit-RED (Cat: 322360, ACD Bio, Newark, CA, USA) as previously described^21^. Sections underwent antigen retrieval in RNAscope target Retrieval buffer (RNAscope Kit) at 95°C for 20 minutes in a decloaking chamber (Biocare, Pacheco, CA, USA), airdried and sections encircled with a hydrophobic pen. Unless stated otherwise slides were washed 2x for a total of 5 minutes. Slides were air dried and treated with Bloxall (Vector Laboratories, SP-6000, Burlingame, CA, USA) for 10 minutes, washed in Milli-Q water and then incubated with protease pre-treatment-3 (RNAscope kit) for 20 minutes at 40°C. Slides were washed in Milli-Q water and incubated with custom-made HIV-1_Bal_ probe (REFL 486631, ACD Bio) for 2 hours at 40°C. Slides were washed in RNAscope Wash Buffer (RNAscope Kit), followed by sequential addition of amplification reagents 1-6 (RNAscope Kit) as follows with washes in RNAscope wash buffer between each amp: Amp 1: 30 minutes 40°, Amp 2: 15 minutes 40°, Amp 3: 30 minutes 40°C, Amp 4: 15 minutes 40°C, Amp 5: 30 minutes room temperature, Amp 6: 15 minutes room temperature. Fast Red chromagen solution was added to sections at a dilution of 1:60 Red-B:Red-A (RNAscope Kit), at room temperature for 5 minutes. Slides were washed firstly in Milli-Q water followed by 2x TBS (Amresco, Cat: 0788) for a total of 5 minutes. Slides were then blocked for 30 minutes (0.1% (w/v) saponin, 1% (w/v) BSA, 10% (v/v) donkey serum) in preparation for immunofluorescence staining. From herein, between each step slides were washed in TBS unless otherwise stated. The first round of primary antibodies were incubated for 1 hour at room temperature, these included goat anti-human langerin (AF2088) and rabbit anti-human CD14 (ab133503). Secondary antibodies were then added for 30 minutes at room temperature, these included donkey anti goat-488 and donkey anti-rabbit-755 (Molecular Probes). Slides were blocked again as above with the addition of 10% (v/v) rabbit serum, followed by 0.5% (v/v) PFA for 15 minutes. Rabbit anti-human CD11c (ab216655) underwent buffer exchange using 50kDa amicon filters (Merck) so as to place antibodies in 0.1M NaHCO3, balanced to a pH of 8.3-8.5. This was followed by conjugation to sulfo-Cyanine5 using the lumiprobe sulfo-Cyanine5 antibody labelling kit (3321-10rxn, Lumiprobe, Hallandale Beach, FL, USA), following the manufacturer’s instructions. The conjugated rabbit CD11c-Cy5 was added to sections and incubated at 4°C overnight. Slides were mounted with prolong SlowFade Diamond Antifade with DAPI (Molecular Probes, Eugene, OR, USA, S36973). Sections were imaged on the Olympus VS120 slidescanner using channels FITC, TRITC, Cy5 and Cy7 using Olympus-asw software (version 2.9) and analysed using Olyvia (version 2.9).

## Results

### Defining human mucosal mononuclear phagocyte subsets by flow cytometry

We have previously optimised protocols for the efficient isolation and definition of MNPs from human skin with minimal surface receptor cleavage and in an immature state to most closely resemble their functional state when they encounter pathogens^40^. We firstly modified our skin flow cytometry gating strategy for use in mucosal tissues (**Figure 1A**). All known tissue MNPs were identified which were; epidermal CD11c^+^ DCs^21^ and LCs and dermal cDC1, cDC2, CD14^+^ autofluorescent macrophages and dermal non-autofluorescent CD14^+^ cells. We also designed the panel to differentiate between non-autofluorescent CD14^+^CD1c^-^ MDMs^49^ and CD14^+^CD1c^+^ MDDCs^31,32,38^. We used this gating strategy to define the relative proportions of each dermal MNP subset in the full range of human anogenital and colorectal tissues that are the actual sites where HIV transmission occurs, as well as abdominal skin for comparison. This included tissues comprised of skin (labia, outer foreskin, glans penis and anal verge), Type II mucosa (vagina, fossa navicularis, ectocervix and anal canal) and type I mucosa (endocervix, penile urethra, rectum and colon) (**Figure 1B**). The relative proportions of cDC1 was relatively consistent across all tissues with the exception of outer foreskin which had slightly higher proportions. In abdominal skin we found that dermal cDC2 were the overwhelmingly predominant cell population in contrast anogenital and colorectal tissues where these cells were present in much smaller proportions. Notably however, a greater proportion of the total cDC2 population expressed langerin in these tissues compared to abdomen (**Figure 1C).** Conversely, CD14-expressing cells were present in higher proportions in anogenital colorectal tissues compared to abdomen and the relative proportions were significantly higher in mucosal tissues compared to anogenital skin (**Figure 1D**). Despite the fact that *in vivo* MDDCs have been predominantly been described as an inflammatory cell we found that they were substantially present across all our uninflamed tissues.

**Figure 1.**
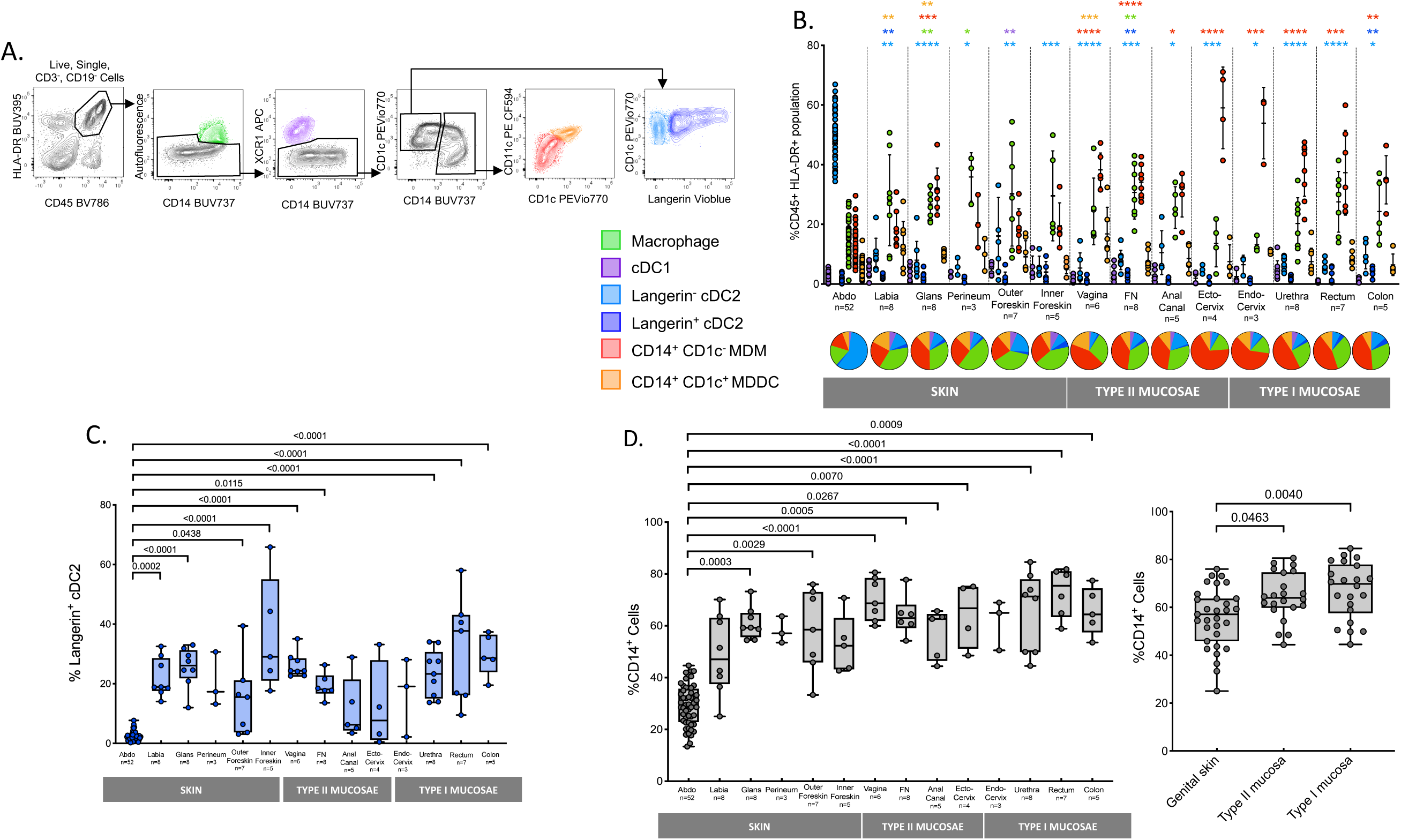
Definition of human dermal/lamina propria mononuclear phagocytes by flow cytometry. **A** Following collagenase digestion six distinct CD3^-^CD19^-^CD45^+^HLA-DR^+^ mononuclear phagocyte subsets from human abdominal skin were defined, (i) tissue resident macrophages (green) were defined as autofluorescent^+^CD14^+^, (ii) type 1 conventional dendritic cells (purple) were defined as autofluorescent^-^, XCR1^+^, CD14^-^, type 2 conventional dendritic cells (blue) were defined as autofluorescent^-^, XCR1^-^, CD14^-^, CD1c^+^, and could be split into two populations, (iii) a langerin^-^ population (light blue) and (iv) a langerin^+^ population (dark blue), (v) CD14^+^CD1c^-^ cells (red) were defined as autofluorescent^-^, XCR1^-^, CD14^+^, CD1c^-^, (vi) CD14^+^CD1c^+^ cells (orange) were defined as autofluorescent^-^, XCR1^-^, CD14^+^, CD1c^+^. Representative plot of *n* = 52 abdominal donors is shown. **B** Relative proportions of each subset of mononuclear phagocyte as a percentage of CD45^+^ HLA-DR^+^ gate across the human anogenital/colorectal tracts were determined and mean +/- standard deviation plotted. Statistics for subsets in each tissue were generated using the Kruskal Wallis test: Dunn’s multiple comparisons, comparing against abdominal skin tissue. *p<0.05; **p<0.01; ***p<0.001; ****p<0.0001. (Abdominal tissue = 52, Labia = 8, Glans Penis = 8, Perineum = 3, Outer Foreskin = 7, Inner Foreskin = 5, Vagina = 6, Fossa Navicularis (FN) = 8, Anal Canal = 5, Ectocervix = 4, Endocervix = 3, Penile Urethra = 8, Rectum = 7, Colon = 5). Underneath, a pie chart for each tissue shows the mean proportion of each dermal/lamina propria subset across the human anogenital/colorectal tracts. **C** The proportion of langerin^+^ cDC2 across the human anogenital/colorectal tissue plotted as box and whisker plots, box representing the upper and lower quartile, central line representing the median, the whiskers the minimum and maximum of each tissue sample and each donor represented by an individual dot.. Statistics and donor numbers as described above for **B. D** The proportion of CD14^+^ mononuclear phagocytes (Macrophages, CD14^+^CD1c^-^ cells, CD14^+^CD1c^+^ cells) across the human anogenital/colorectal tract, left: looking at individual anogenital/colorectal tissue, right: pooled data for genital skin (Labia, Glans penis, Perineum, Outer Foreskin, Inner Foreskin), Type II mucosae (Vagina, Fossa Navicularis, Anal Canal, Ectocervix) and Type I mucosae (Endocervix, Penile Urethra, Rectum, Colon) graphed as described above for **C**. Statistics and donor numbers as described above for **B**.

### Transcriptional profiling

As CD14^+^ cells have been shown to play an important role in HIV transmission we firstly focused our study on these cells. As we found both CD14^+^CD1c^+^ MDDC and CD14^-^CD1c^-^ MDM in all our uninflamed tissue samples we carried out RNAseq analysis to compare the transcription profile of these cells to other tissue MNPs; LCs, dermal langerin^-^ cDC2 and their langerin-expressing counterparts (**Figure 2A**). As expected, LCs were the most distinct population and, in agreement with the literature, cDC2 clustered together regardless of langerin expression^50^ and CD14^+^CD1c^-^ cells were transcriptionally very similar to autofluorescent macrophages^36^ whereas CD14^+^CD1c^+^ cells aligned more closely with DCs^38,39^. Comparison of *ex vivo* MDM cells with *ex vivo* MDDC cells showed 501 differentially expressed genes. CD14^+^CD1c^+^ MDDCs expressed lower levels of the lentiviral restriction factor APOBEC3G which is a known inhibitor of HIV-1 infectivity^51^ (**Figure 2B).**

**Figure 2.**
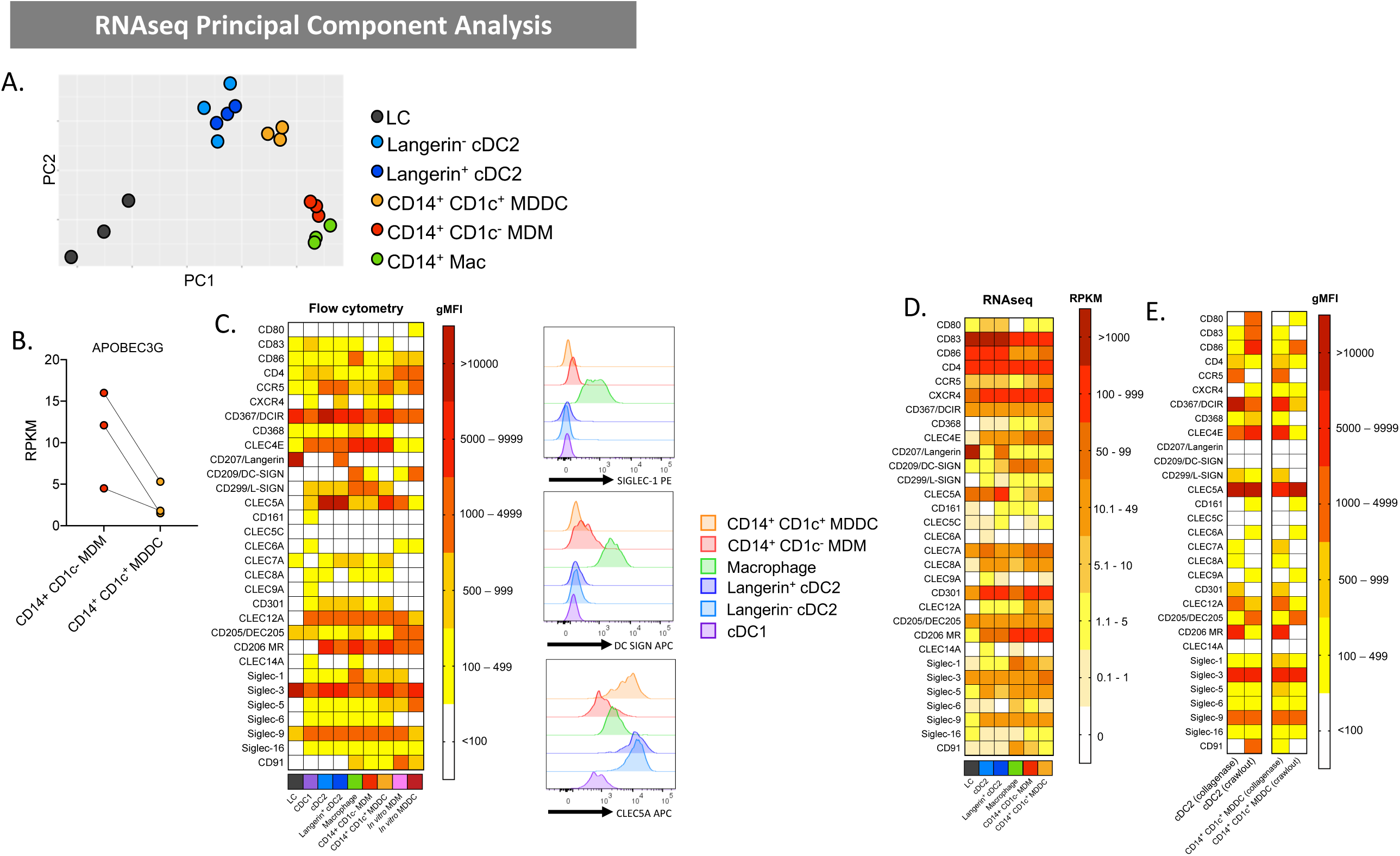
Genotypic and phenotypic profiling of epidermal and dermal human abdominal skin mononuclear phagocytes. **A** Dermal macrophages, cDC2 (langerin^-^ and langerin^+^), CD14^+^CD1c^-^ cells and CD14^+^CD1c^-^ cells as well as epidermal Langerhans cells were liberated from *n* = 3 abdominal skin donors and FACS isolated for RNAsequencing (cDC1 were omitted from analysis due to low donor numbers). A PCA plot shows clustering of cell subsets. **B** HIV restriction factor APOBEC3G gene expression by RNAseq for three donors for CD14^+^CD1c^-^ and CD14^+^CD1c^+^ cells. Reads Per Kilobase of transcript, per Million mapped reads (RPKM) is plotted with each dot representing an individual donor and lines matching donors. **C** Epidermal LCs and dermal MNPs were liberated from tissue using enzymatic digestions and surface receptor expression evaluated by flow cytometry and compared to *in vitro* Monocyte-derived macrophages (MDM) and Monocyte-derived dendritic cells (MDDC). Left: The geometric mean fluorescent intensity (gMFI) minus the isotype for each subset was calculated and mean plotted in a heat map (n= 2-11). Right: Representative histograms of one donor for cell surface expression of three c-type lectin receptors of interest, Siglec-1, DC-SIGN, CLEC5a. **D** A heatmap was also generated from the RNAseq data to investigate the expression of the corresponding genes investigated by flow cytometry in **C. E** Flow cytometry analysis shows differences in cell surface expression on immature (enzymatically digested-derived cells) vs mature (spontaneous migration-derived cells) for left: cDC2 and right: CD14^+^CD1c^+^ cells.

### Determination of surface receptor expression on skin mononuclear phagocytes

Previously we have carried out gene expression studies to examine CLR expression by *ex vivo* derived skin DC subsets derived by collagenase digestion and compared them to model *in vitro* derived cells^52^. However we did not include cDC1 or autofluorescent macrophages nor did we divide the non-autofluorescent CD14-expressing cells according to CD1c expression or cDC2 by langerin expression. Importantly, we also did not examine the surface expression of these proteins. We therefore used our flow cytometry gating strategy (**Figure 1A)** to determine the surface expression levels of a wide range of surface molecules including HIV entry receptors, costimulatory molecules and lectin receptors involved in pathogen recognition (CLRs and Siglecs) on each MNP subset obtained via enzymatic digestion (for immature cells) or spontaneous migration (for mature cells). We also included MDDC and MDM derived *in vitro* from blood CD14 monocytes which are commonly used as model cells (**Figure 2C**). Importantly, we used appropriate collagenase enzyme blends which we have previously determined do not cleave each specific surface receptor^40^. As we had carried out RNAseq for cells derived by enzymatic digestion we were able to compare gene and surface expression profiles for immature cells isolated by this method and in almost all cases similar trends were observed (**Figure 2D**).

All MNPs expressed the key HIV entry receptor CD4 but cDC1 and LCs expressed very low levels of the HIV entry co-receptor CCR5. These two subsets also differed from other MNPs subsets in that they expressed far fewer lectin receptors. LCs only expressed langerin, DCIR, DEC205, Siglec-3 and Siglec-9 as well as very low levels of CLEC7A, CLEC4D and CLEC4E. cDC1 expressed slightly more lectin receptors than LCs but generally at lower levels than other MNPs and uniquely expressed CLEC9A consistent with the literature. Other than langerin expression, we did not identify any differences in surface expression molecules between langerin-expressing and non-expressing cDC2.

#### DC-SIGN, Siglec-1 and CLEC5A differentiate DC from macrophage like cells

Interestingly, the HIV binding CLR DC-SIGN, previously thought to be a DC marker and shown to be involved in capture of HIV and transfer to T cells^11,32-34,49-52^, was not expressed by any *ex vivo* DC subsets or CD14^+^CD1c^+^ MDDCs (although gene expression was detected). It was expressed highly by *ex vivo* autofluorescent macrophages and *in vitro* derived MDDCs and at low levels by both *ex vivo* CD14^+^CD1c^-^ and *in vitro* MDMs (**Figure 2C and D).** Similarly Siglec-1, a lectin shown to be important in DC mediated HIV uptake^15,53,54^ was expressed most highly by autofluorescent macrophages and *ex vivo* MDMs but at lower levels on *ex vivo* MDDCs lower still on cDC1 and cDC2 and not all by *in vitro* derived MDDCs (**Figure 2C and D)**. Finally, CLEC5A was expressed very highly by cDC2 and *ex vivo* MDDCs but at lower levels on other CD14^+^ cells. Other than this, all CD14^+^ *ex vivo* derived cells showed very similar surface expression profiles. Taken together the expression profiles of DC-SIGN, Siglec-1 and CLEC5A add further weight to the hypothesis that CD14^+^CD1c^+^ cells are DC-like and CD14^+^CD1c^-^ cells are macrophage like.

#### Differences between model cells and ex vivo derived cells

In addition to what is described above, model *in vitro* derived MDDCs and MDM differed from *bona fide ex vivo* derived cells in several important ways; similar to LCs, they did not express L-SIGN, CLEC7A, CLEC10A and Siglec-6 but, unlike LCs did not express CLEC4D or langerin. They also expressed much higher levels of CD4 and DEC205 than any other cell type.

#### Differential expression of surface receptors between immature and mature cells

Using the spontaneous migration method we were only able to reliably isolate langerin^-^ cDC2 and CD14^+^CD1c^+^ MDDCs as described previously^40^ (**Figure 2E)**. As expected cells isolated by this method expressed much higher levels of CD80 and CD86 than cells isolated by enzymatic digestion confirming their mature state. Interestingly cDC2, but not MDDCs expressed CD83. Cells isolated by spontaneous migration expressed much lower (or none) of the following molecules: CLEC4A, CLEC4K (langerin), CLEC4L (DC-SIGN), CLEC8A, CLEC10A, CLEC12A and Siglec-1, while CLEC4E was down regulated by MDDCs but not cDC2. Conversely CLEC13B (DEC205) was upregulated. In terms of HIV entry receptors all cell subsets expressed CD4 regardless of their method of isolation. As we showed previously, CCR5 expression was not detected on immature LCs^21^ or cDC1 or any mature cells. CXCR4 was expressed at very low levels or not at all by mature cells but was expressed at slightly higher levels by LCs and MDDCs isolated by the spontaneous migration method.

### CD14^+^ dermal cell morphology and tissue residency

As DCs and macrophages are morphologically different^27^ we next compared the morphology of dermal MDDCs, MDMs and cDC2 using Giemsa staining (**Figure 3A**). Consistent with the transcriptional profiling and surface expression profile, we found that MDMs looked like macrophages with a ‘fried egg’ like appearance and containing many large intracellular vacuoles whereas MDDCs cells were morphologically more similar to DCs with none or few vacuoles present.

**Figure 3.**
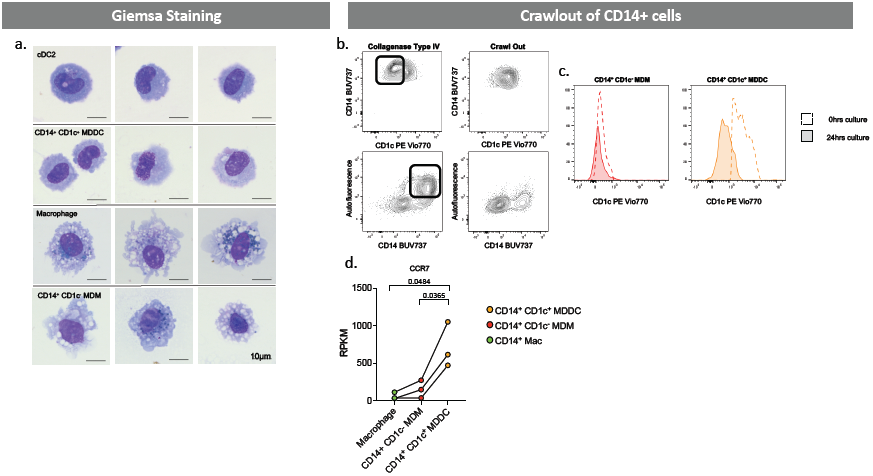
Morphological and migratory properties of CD14-expressing human tissue mononuclear phagocytes. **A** Human dermal MNPs were isolated from abdominal skin by collagenase digestion, with each defined subset FACS isolated and Giemsa stained to investigate morphology. **B** The migratory capabilities of CD14-expressing cells (macrophages, CD14^+^CD1c^-^ cells, CD14^+^CD1c^-^ cells). From a single donor dermal sheets were cultured for 24 hours and supernatant collected for cells which had undergone spontaneous migration (crawl out), tissue was then collagenase digested to liberate cells which did not undergo spontaneous migration. Contour plots show the phenotype of cells which crawled out compared to those which remained within the tissue, box highlighting cells which remained in tissue. **C** From a separate donor CD14^+^CD1c^-^ cells and CD14^+^CD1c^+^ cells were FACS isolated following collagenase digestion and cultured for 0 hours (unfilled, dotted histogram) and 24 hours (filled, continuous histogram) to determine the difference in CD1c expression on these cells before and after maturation. **D** Gene expression of chemokine receptor CCR7 determined by RNAseq for CD14-expressing MNPs from three abdominal skin donors. Reads Per Kilobase of transcript, per Million mapped reads (RPKM) is plotted, with each donor represented by an individual dot and lines connecting donors. Statistics were generated using a paired T -test.

DC and macrophages also differ in tissue residency which has important implications for transmission of HIV to CD4 T cells; DCs migrate out of tissue to lymph nodes which are rich in CD4 T cells whereas macrophages remain tissue resident. Consistent with this migration differential, DCs can be isolated from tissues by the spontaneous migration method (albeit in an activated state)^40^ whereas macrophage isolation requires enzymatic digestion. We therefore compared the ability of dermal MDDCs and MDMs to migrate out of tissue spontaneously (**Figure 3B)**. Consistent with their DC-like transcriptional and morphological phenotype MDDCs migrated out of tissue whereas MDMs did not, consistent with their macrophage like phenotype. To confirm that the MDMs were still present within the tissue, we digested the tissue after spontaneous migration and found that they were still present. We next sorted MDM and MDDCs after isolation by enzymatic digestion and confirmed that MDMs did not upregulate CD1c with culture, whereas MDDCs showed a slight downregulation of CD1c, suggesting the population of cells which migrated out of tissue were *bona fide* CD14^+^CD1c^+^ MDDCs. **(Figure 3C)**. Finally, as CCR7 is a key chemokine receptor required for DC migration out of tissue and shown in lung to be expressed by MDDC^39^, we compared the CCR7 gene expression levels in both cells types and showed that CCR7 was expressed much more highly on MDDCs (**Figure 3D**). Taken together this shows that MDDCs migrate out of tissue whereas MDMs do not.

### CD14^+^ dermal cell MNPs take up HIV via Siglec-1

We next compared the ability of *ex vivo* derived CD14-expressing cell subsets to take up HIV after 2 hours of exposure using both the lab-adapted Bal strain and the transmitted/founder Z3678M strain^21,55^. All subsets efficiently took up the virus but MDDCs took up significantly more **(Figure 4A)**. We were also able to observe both MDDCs and MDMs interacting with HIV *in situ* in inner foreskin using our RNAscope HIV detection assay^21^ indicating that HIV can penetrate into the sub-epithelium and interact with these cells *in situ* (**Figure 4B)**. CD14-expressing cells have previously been shown to play a role in HIV transmission and we and others have shown that DC-SIGN is a key HIV binding receptor expressed on these cells and is necessary for uptake and transfer to CD4 T cells^11,33-35,56-59^. Furthermore we have shown blocking the ability of HIV to bind both DC-SIGN and mannose receptor on MDDCs derived *in vitro* from CD14^+^ monocytes significantly blocks HIV uptake^11,12^. However, we observed here that another key HIV binding lectin receptor, Siglec-1^15,53,54,60^, was expressed by all CD14^+^ sub-epithelial cells and at much higher levels than CD14^-^ MNP subsets (**Figure 2C).** We therefore examined Siglec-1 expression on HIV infected and by stander cells derived from labia, inner foreskin and colon. In all CD14-expressing cell types we found that cells that had taken up HIV expressed significantly higher levels of Siglec-1 than cells that did not take up HIV, indicating that Siglec-1 expression correlates with HIV uptake (**Figure 4C)**. Furthermore, in all specimens Siglec-1 was expressed most highly in autofluorescent tissue resident macrophages followed by MDM then MDDCs. This finding was confirmed further in colonic tissue (**Figure 4D**). Using cells derived from labia, foreskin and colon we showed that a Siglec-1 antibody was able to partially block HIV uptake further implying a role of Siglec-1 in HIV uptake by CD14-expressing cells (**Figure 4E).** We also measured the ability of all subsets to transfer HIV to CD4 T cells at 2 hours (**Figure 4F**). We found no statistically significant difference between the three subsets’ ability to transfer the virus, although autofluorescent tissue resident macrophages (which expressed the highest levels of Siglec-1) were able to transfer the virus much more efficiently in 2 of the 4 donors. Therefore, Siglec-1 expression does not explain why *in vivo* MDDCs cells take up HIV more efficiently than other CD14-expressing cells.

**Figure 4.**
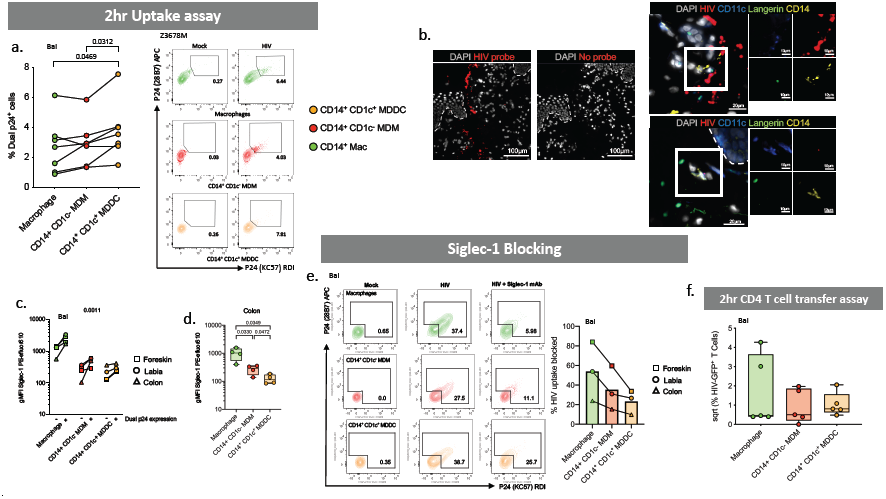
HIV uptake and transfer capacity of dermal CD14^-^expressing mononuclear phagocytes. **A** MNPs were liberated from human abdominal skin using and CD45^+^HLA-DR^+^ live cells FACS isolated. Mixed dermal populations were incubated for 2 hours with HIV_Bal_ or HIV_Z3678M_ or mock treated, thoroughly washed and stained for surface markers and two antibody clones to HIV p24 (KC57 and 28B7) for flow cytometry analysis to determine percentage dual p24^+^ cells. Left: HIV_Bal_ results graphed with each donor represented by individual dots, donor matched by connected lines (n=7). Statistics were generated using Wilcoxon test. Right: Representative contour plots for HIV_Z3678M_ shown, gating on dual p24^+^ cells for each subset. **B** Human inner foreskin explants were treated with HIV for 2-3 hours before being fixed and paraffin embedded. After sectioning, HIV RNA was visualized using RNAscope and immunofluorescent staining for langerin, CD11c, CD14 and DAPI. Left: Representative images of one donor, with and without the addition of the HIV specific RNAscope probe. Right: Macrophages and CD14^+^CD1c^-^ cells could not be individually distinguished from each other but could be visualised as a group (top) while CD14^+^CD1c^+^ cells could also be visualised (bottom). **C** Human MNPs from anogenital/colorectal tissue (2x foreskin: squares, 1x labia: circle, 1x colon: triangle), were treated as in **A**. The expression of Siglec-1 was assessed by geometric mean fluorescence intensity (gMFI) minus matching FMO control, on dual p24^+^ cells vs p24^-^ cells. Each dot represents an individual donor with lines matching the same donor for each subset. Statistics measured the difference in p24^+^ cells versus p24^-^ cells using a three-way ANOVA. **D** Siglec-1 expression on CD14-expressing subsets was determined in *n* = 4 colon donors. **E** Mixed dermal MNPs were treated as above however before incubation with HIV, cells were pretreated with Siglec-1 mAb for 30 minutes. Left: representative plot of HIV uptake assessed by dual p24^+^ cells in Mock, HIV treated and HIV + Siglec-1 mAb samples from foreskin donor. Right: Percentage of HIV uptake that was blocked using Siglec-1 mAb across three donors of anogenital/colorectal tissue (Foreskin: square, Labia: circle, Colon: Triangle), with each point representing individual donors matched with joining lines, and column representing mean. **F** Sorted CD14-expressing dermal cells were incubated with HIV_Bal_ for 2 hours and then thoroughly washed off. JLTRs cells were added to MNPs at a ratio of 4:1 and cultured for a further 96hrs in human fibroblast conditioned media. Transfer of HIV to T cells was assessed using flow cytometry to plot the square root percent of GFP^+^ T cells, with each dot representing an individual donor (n=5).

### MDDCs are most susceptible to HIV infection

We next investigated the ability of dermal CD14-expressing cells to become productively infected with HIV and to transfer the virus to CD4 T cells at later time points. As the expression of the HIV entry receptors CD4 and CCR5 are essential for productive HIV infection, we firstly measured the surface expression of these molecules on *ex vivo* cells abdominal skin and found that MDDCs expressed higher levels of CCR5 than other CD14-expressing populations (**Figure 5A)**. We then confirmed this observation in cells derived from colon tissue which is more relevant to HIV transmission (**Figure 5B)**. Corresponding with their higher CCR5 surface expression, we also found that supernatants derived from infected abdominal MDDCs cultures were able to infect greater numbers of CD4 TZMBL cells than other CD14-expressing subsets **(Figure 5C)**. Despite using our optimised *ex vivo* derived skin MNP culture methods^40^, we were unable to measure direct infection of CD14-expressing cell subsets by flow cytometry (as we have done previously with other *ex vivo* derived skin MNPs^13,21^) as only small cell yields could be derived for each specific cell subsets (particularly MDDCs), even from very large pieces of abdominal skin, and too few live cells could be detected after 96 hours of culture. Corresponding with their higher levels of HIV infection, we finally showed that MDDCs were the most efficient CD14-expressing cells at transferring HIV to CD4 T cells (**Figure 5D).**

**Figure 5.**
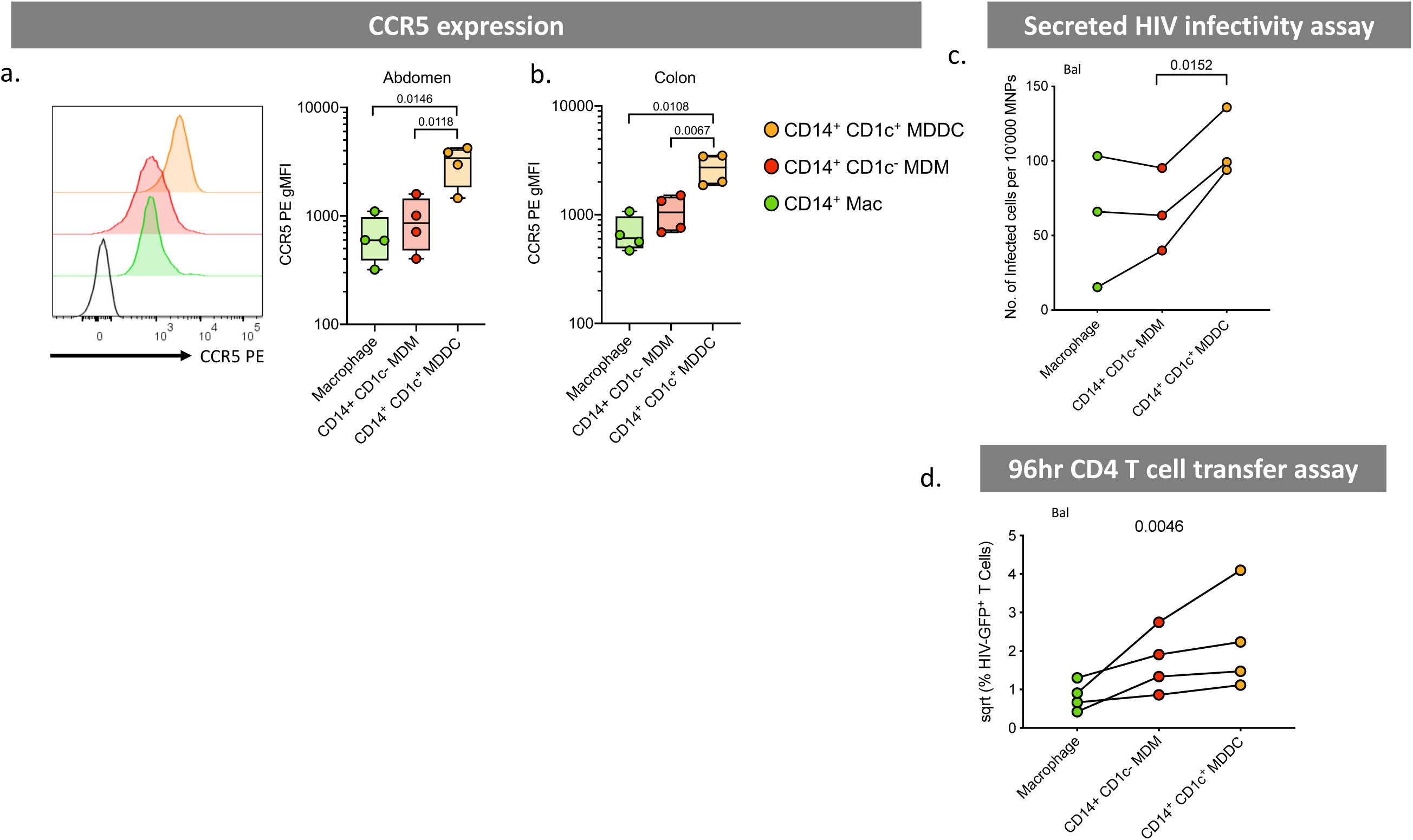
HIV infectability and transfer capacity of CD14-expressing dermal mononuclear phagocytes. **A** CCR5 expression was determined on CD14-expressing MNPs liberated from collagenase type IV digested tissue. Left: Histogram representing one abdominal skin donor with black, unfilled histogram representing FMO control. Right: geometric mean fluorescent intensity (gMFI) minus FMO control plotted from four abdominal skin donors, graphed as box and whisker plots, box representing the upper and lower quartile, central line representing the median, the whiskers the minimum and maximum of each cell subset and each donor represented by an individual dot. Statistics were generated using a Paired T-test. **B** CCR5 expression was investigated in four colon donors as presented as in A. **C** CD14-expressing MNP subsets were FACS isolated and incubated with HIV_bal_ for 2 hours before being thoroughly washed off. Cells were then cultured for an additional 96 hours in human skin fibroblast conditioned media. Cell supernatants were taken and assessed for secreted HIV using a TZMBL infection assay, with the number of infected cells per 10,000 MNPs calculated and graphed. Individual dots represent three donors matched by connecting lines. Statistics were generated as in A. **D** JLTRs were added to cell cultures from **C** after supernatants removed, at a ratio of 4:1 and cultured for a further 96 hours. Transfer of HIV to T cells was determined using flow cytometry to assess the percent of GFP^+^ JLTRs. Raw data was square root normalised and plotted, with each dot representing four individual donors with connecting lines matching donors. Statistics were generated using a Friedman test.

### Langerin^+^ cDC2 are enriched in anogenital tissues and the most efficient cells at HIV uptake and infection

We next turned our attention to sub-epithelial cDC2 which have been understudied in HIV transmission. We have recently shown that these cells exist in the epidermis of human anogenital cutaneous tissues where they preferentially interact with HIV and transmit it to T cells^21^. Interestingly, we found that, similar to epidermis, langerin^+^ sub-epithelial cDC2 were significantly enriched in anogenital tissues compared to abdominal skin **(Figure 1C)**. We also found that within 2 hours these cells took up much more HIV than their langerin^-^ counterparts and cDC1 using both a lab-adapted and transmitted founder HIV strain (**Figure 6A)** and could be visualised interacting with the virus *in situ* using RNAscope following topical infection of foreskin explants (**Figure 6B)**. Correspondingly, they also transferred HIV to CD4 cells more efficiently at 2 hours than langerin^-^ cDC2 (**Figure 6C).** Interestingly, although langerin^+^ cDC2 did not express higher levels of CCR5 (**Figure 6D**) or CD4 (**Figure 6E)** than their langerin^-^ counterparts, they nevertheless produced higher levels of both HIV lab adapted BaL strain and a transmitted funder strain after 96 hours indicating they supported higher levels of infection **(Figure 6F).** We attempted to block HIV uptake and transfer to CD4 cells using a langerin antibody, as we have done previously with LCs^13^, and also to carry out a 96 hour MNP-T cell transfer assay but unfortunately these experiments proved impossible due to very small numbers of langerin expressing sub-epithelial cells that are able to be extracted from tissue.

**Figure 6.**
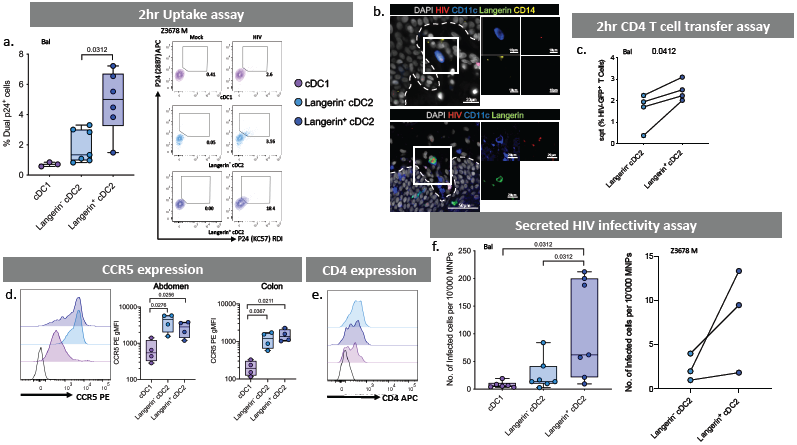
Uptake, infectivity and transfer capacity of dermal dendritic cells. **A** Enzymatically-liberated cells from abdominal skin were FACS isolated to obtain a CD45^+^HLA-DR^+^ population and cultured with HIV_Bal_ or HIV_Z3678M_ or Mock treated for 2 hours before being washed off three times in PBS. Cells were stained for surface markers and two p24 clones, before analysis by flow cytometry. Left: HIV_Bal_ treated cells percentage of dual p24^+^ cells plotted as box and whisker plots, box representing the upper and lower quartile, central line representing the median, the whiskers the minimum and maximum of each cell subset and each donor represented by an individual dot (n=7, n=3 for cDC1 due to low cell numbers). Statistics were generated using a Wilcoxon test. Right: representative contour plots for HIV_Z3678M_ gating on dual p24^+^ cells. **B** Human inner foreskin explants were treated with HIV for 2-3 hours, before being fixed, paraffin embedded and sectioned. HIV was visualised using RNAscope alongside immunofluoerescence for CD11c, CD14 and langerin as well as DAPI to identify Langerin^-^ cDC2 (top) and langerin^+^ cDC2 (bottom). **C** FACS isolated cDC2 divided by langerin expression were cultured with HIV_Bal_ for 2 hours before being washed off three times. JLTR cells (CD4 T cells with GFP under the HIV promoter) were then co-cultured for 96 hours in fibroblast conditioned media. Flow cytometry was used to analyse the percent of GFP^+^ T cells. Data was square root normalised and graphed with each donor represented by an individual dot and lines connecting matched donors (n=4). Statistics were generated using a paired T test. **D** HIV co-receptor CCR5 expression was analysed by flow cytometry. Left: representative histogram of one donor, black unfilled histogram represents FMO control. CCR5 gMFI minus the FMO for four abdominal skin donors (middle) and four colon donors (right) plotted as box and whisker plots, as in A. Statistics were generated using paired T-tests. **E** HIV receptor CD4 expression levels on one abdominal skin donor with isotype control represented by black, unfilled histogram. **F** Cell supernatants from FACS isolated dermal DCs infected with HIV were taken for TZMBL infectivity assays and the number of infected TZMBL cells per 10^4^ MNPs were calculated. Left: HIV_Bal_ infectivity plotted as box and whisker plots as in **A**, n=7. Statistics were generated using a Wilcoxon test. Right: HIV_Z3678M_ infectivity assay with each dot representing individual donors matched by connecting lines (n=3). Due to low cell numbers cDC1 data could not be obtained.

## Discussion

In this study we have investigated the role that sub-epithelial MNPs play in HIV transmission. We began by thoroughly defining the relative proportions MNPs that are present in all human anogenital and colorectal tissues which are the sites of HIV transmission. In doing so we revealed that CD14-expressing cells predominate in these tissues in contrast to abdominal skin. CD14-expressing cells were found in greater proportions in internal mucosal tissues compared to external genital skin. We identified two MNPs subsets that may play a dominant role in HIV transmission; CD14^+^CD1c^+^ MDDCs and langerin^+^ cDC2, both of which were present in higher proportions in human anogenital and colorectal tissues compared to abdominal skin.

Tissue macrophages express CD14 and have been classically defined in tissue by their high levels of autofluorescence^27,61^. Non-autofluorescent tissue CD14^+^ cells have traditionally been referred to as CD14 DCs which have been well characterised in their ability to bind to and capture HIV and transmit it to CD4 T cells in model systems^11,17,43,62-64^ as well as in intestinal tissue^33-35^ and more recently in cervical tissue^15^. The ability of these cells to transmit the virus to CD4 T cells has been assumed to be associated with the potent antigen presenting function of DCs. However, transcriptional profiling has recently led to non-autofluorescent CD14^+^ CD1c^-^ cells in skin to be redefined as MDMs^49^. This is puzzling as macrophages are thought to be weak antigen presenting cells to naïve T cells, especially when compared to DCs and they do not migrate out of tissue to lymph nodes^61^. As CD14^+^CD1c^+^ MDDCs have been described in inflamed tissue^31,32,38^ we therefore carried out transcriptional profiling and found that, in agreement with the literature, CD14^+^CD1c^-^ cells transcriptionally aligned very closely with macrophages^49^ and that CD14^+^CD1c^+^ cells aligned more closely with DCs^38^. Adding weight to the hypothesis that these cells were DC-like, we showed that CD14^+^CD1c^+^ cells morphologically resembled DCs and, expressed *CCR7* and spontaneously migrated out of tissue as DCs are known to do, unlike CD14^+^CD1c^-^ cells which morphologically resembled macrophages, did not express *CCR7* and remained tissue resident. These results support the literature which shows that CD1c is an essential marker to define cDC2^65^ and inflammatory MDDCs have been shown to express CD14 and CD1c in patients with rheumatoid arthritis and cancer^32,38^. Indeed, they recently showed that all human CD1c^+^ inflammatory DCs derived from ascites are monocyte-derived cells and therefore not cDC1 or cDC2 which are derived from specific bone marrow precursors^31^. However, our findings here are novel as these cells have not been previously defined in healthy human anogenital tissue which form the portals of HIV entry and we show that they did differ from inflammatory MDDCs in that they did not express CD1a, CD141 or DC-SIGN.

We next investigated the way CD14-expressing cells interacted with HIV-1_BaL_ and a transmitted/founder clinical HIV-1 strain and found that MDDCs cells took up significantly more HIV within 2 hours than MDMs and autofluorescent macrophages. They also expressed higher levels of the HIV entry receptor CCR5 and correspondingly supported higher levels of productive infection resulting in higher levels of infectious virion secretion and higher levels of transfer of the virus to CD4 T cells. They also expressed lower levels of the HIV restriction factor *APOBEC3G* which could further account for the higher levels of secreted virions by these cells. Uptake of HIV by MNPs at early time points is mediated by lectin receptors and previously DC-SIGN has shown to be key a lectin receptor involved in HIV uptake by CD14-expressing cells and has also been implicated in transfer to CD4 T cells^11,30,33,34,58^. Similarly Siglec-1 has also been implicated^15,60^. However, MDDCs did not express DC-SIGN so this receptor cannot be responsible for the efficient early uptake observed at 2 hours by these cells. This is an important observation as soluble DC-SIGN designed to block HIV interacting with MNPs has been proven ineffective in blocking HIV transmission^66^. MDDCs did however express Siglec-1, but at lower levels than MDMs and autofluorescent macrophages which expressed the highest Siglec-1 levels. We therefore investigated the role of Siglec-1 in early uptake and found that in all CD14-expressing cell types, those cells that contained HIV expressed higher levels of Siglec-1 than uninfected cells. Furthermore, a Siglec-1 blocking antibody was able to partially block HIV uptake which corresponded with the levels of Siglec-1 surface expression. Therefore, the mechanism which allows MDDCs to take up HIV more efficiently than other CD14-expressing cells remains to be elucidated and will be the subject of a future study. However, other than DC-SIGN did not detect any difference in the expression profiles of **<u>known</u>** HIV binding CLRs between MDDCs and other CD14^+^ cells. MDDCs did however express higher levels of CLEC5A which opens an avenue of investigation.

Despite their importance in antigen presentation, cDC2 have been understudied in HIV transmission with almost all studies focussing on langerin-expressing LCs or DC-SIGN expressing CD14^+^ cells^10,67^. Recently we reported that cDC2-like cells were present in the epidermis of anogenital tissues where they predominated over LCs and preferentially become infected with HIV and transmitted the virus to CD4 T cells^21^. Pena-Cruz and colleagues made similar observations in vaginal epithelium^22^. Importantly, we showed that in anogenital tissues the majority of these cells expressed langerin. We therefore investigated the role of lamina propria cDC2 in HIV transmission. We noticed that a greater proportion of sub-epithelial cDC2 in anogenital and colorectal tissues expressed langerin than in abdominal dermis. This was especially the case in the inner foreskin, penile urethra, vagina and rectum. Importantly, we found that langerin-expressing cDC2 were much more efficient at HIV uptake after 2 hours than their non-langerin-expressing counterparts and were correspondingly much more efficient at transferring the virus to CD4 T cells at the same early time point. Furthermore, despite the fact that no differences in surface CCR5 expression were detected between the two cells types, langerin-expressing cDC2 also secreted higher levels of infectious HIV virions after 96 hours using both the lab-adapted Bal strain and a clinical transmitted/founder strain. Other than langerin expression these two cell types expressed an identical array of surface lectin receptors so we hypothesise that langerin must be mediating these effects. In LCs langerin is well known to bind HIV and mediate efficient HIV uptake^13,67-69^ so we hypothesise here that these cells efficiently bind HIV via langerin which concentrates HIV on the cells surface allowing for greater HIV-CCR5 binding and infection as well as endocytic uptake. We made multiple attempts to test this hypothesis using a langerin blocking antibody as we did previously with LCs^13^ but unfortunately these experiments proved impossible using the very small numbers of this cell type we could extract from abdominal or genital skin or mucosal tissue.

A clear strength of this study is that it has been exclusively conducted using human tissues including all the anogenital and colorectal tissues that HIV may encounter during sexual transmission. However, these kinds of experiments also come with limitations. They are very laborious and time consuming and only a very small number of cells can be isolated for each specific subset, especially MDDCs and even more so for langerin^+^ cDC2 which were both a key focus of this study. This severely limits the parameters that can be included in our assays and many experiments proved impossible such as blocking assays using langerin^+^ cDC2. Furthermore, these cells are very difficult to culture once isolated from tissue and despite our optimised cell extraction and *ex vivo* culture protocols^40^ functional assays requiring 96 hours of culture were extremely difficult to perform. We did manage to perform HIV infection assays using infected MNP culture supernatants but there were too few live cells to gate on to measure direct infection of any MNP cell type by flow cytometry as we have done previously for epidermal cells^13,21^ and we were unable to perform transfer assays at 96 hours for langerin^+^ cDC2. These constraints also meant that we could only repeat a few key observations with a clinical transmitted/founder strain. All uptake assays were confirmed at least once using this strain and also the cDC2 infectivity assays. In all experiments similar trends were observed with both virus strains. Furthermore, we have previously observed similar trends using the same two strains of HIV in epidermal LCs and CD11c^+^ DCs^21^.

In conclusion, we have identified two sub-epithelial MNPs that may play a role in transmission of HIV; CD14^+^CD1c^+^ MDDCs and langerin^+^ cDC2. Both were able to preferentially take up the virus within 2 hours and support higher levels of HIV infection than other MNP subsets. Previously, many studies have focussed on the role of LCs in sexual transmission of HIV ^67^ as these were considered most likely to interact with HIV as they are closest to the epithelial surface. We also demonstrated the importance of a second epidermal DC subset, resembling activated cDC2^21^. However, as genital trauma and inflammation are clearly associated with HIV transmission, especially in sub-Saharan Africa^3-5^, and current PrEP regimens, if available, are not efficient at blocking transmission across an inflamed mucosa^7-9^, it is important that the role of sub-epithelial MNPs are also examined as we have done here. This is relevant to vaccine design as, to protect against initial infection, the route of transmission and which local immune defence mechanisms can be harnessed or impaired needs to be understood. The current targets of systemic vaccines as broadly neutralising antibodies, antibody dependent cytotoxicity and systemic CD8 T cells may not be enough. Local immunity in the anogenital mucosa such as resident memory T cells could be induced by mucosal vaccines or maintained by local tissue DCs after initial stimulation by systemic vaccine adjuvants in lymph nodes draining the site of application. Therefore, it is important to determine which specific subsets of MNPs pick up the virus and deliver it to specific subsets of CD4 T cells. Here we have identified two MNP subsets of interest and together with our recently discovered epidermal CD11c^+^ DCs^21^ and LCs, the next step is to determine with which T cells subsets these cells interact. Finally, defining the interactions between HIV and its mucosal target cells should aid in the development of blocking agents that can be used in modified PrEP regimens to block MNP infection via the anogenital and colorectal mucosa.

## References

1. de Visser, R.O., et al. Change and stasis in sexual health and relationships: comparisons between the First and Second Australian Studies of Health and Relationships. Sex Health 11, 505–509 (2014).

2. Kojima, N., Davey, D.J. & Klausner, J.D. Pre-exposure prophylaxis for HIV infection and new sexually transmitted infections among men who have sex with men. AIDS 30, 2251–2252 (2016).

3. Esra, R.T., et al. Does HIV Exploit the Inflammatory Milieu of the Male Genital Tract for Successful Infection? Front Immunol 7, 245 (2016).

4. Passmore, J.A., Jaspan, H.B. & Masson, L. Genital inflammation, immune activation and risk of sexual HIV acquisition. Curr Opin HIV AIDS 11, 156–162 (2016).

5. Patel, P., et al. Estimating per-act HIV transmission risk: a systematic review. AIDS 28, 1509–1519 (2014).

6. Powers, K.A., Poole, C., Pettifor, A.E. & Cohen, M.S. Rethinking the heterosexual infectivity of HIV-1: a systematic review and meta-analysis. Lancet Infect Dis 8, 553–563 (2008).

7. Abdool Karim, S.S., Passmore, J.S. & Baxter, C. The microbiome and HIV prevention strategies in women. Curr Opin HIV AIDS 13, 81–87 (2018).

8. Klatt, N.R., et al. Vaginal bacteria modify HIV tenofovir microbicide efficacy in African women. Science 356, 938–945 (2017).

9. McKinnon, L.R., et al. Genital inflammation undermines the effectiveness of tenofovir gel in preventing HIV acquisition in women. Nat Med 24, 491–496 (2018).

10. Harman, A.N., Kim, M., Nasr, N., Sandgren, K.J. & Cameron, P.U. Tissue dendritic cells as portals for HIV entry. Rev Med Virol 23, 319–333 (2013).

11. Bernhard, O.K., Lai, J., Wilkinson, J., Sheil, M.M. & Cunningham, A.L. Proteomic analysis of DC-SIGN on dendritic cells detects tetramers required for ligand binding but no association with CD4. J Biol Chem 279, 51828–51835 (2004).

12. Lai, J., et al. Oligomerization of the macrophage mannose receptor enhances gp120-mediated binding of HIV-1. J Biol Chem 284, 11027–11038 (2009).

13. Nasr, N., et al. Inhibition of two temporal phases of HIV-1 transfer from primary Langerhans cells to T cells: the role of langerin. J Immunol 193, 2554–2564 (2014).

14. Mikulak, J., Di Vito, C., Zaghi, E. & Mavilio, D. Host Immune Responses in HIV-1 Infection: The Emerging Pathogenic Role of Siglecs and Their Clinical Correlates. Front Immunol 8, 314 (2017).

15. Perez-Zsolt, D., et al. Dendritic Cells From the Cervical Mucosa Capture and Transfer HIV-1 via Siglec-1. Front Immunol 10, 825 (2019).

16. Menager, M.M. & Littman, D.R. Actin Dynamics Regulates Dendritic Cell-Mediated Transfer of HIV-1 to T Cells. Cell 164, 695–709 (2016).

17. Turville, S.G., et al. Immunodeficiency virus uptake, turnover, and 2-phase transfer in human dendritic cells. Blood 103, 2170–2179 (2004).

18. Ganor, Y., et al. Within 1 h, HIV-1 uses viral synapses to enter efficiently the inner, but not outer, foreskin mucosa and engages Langerhans-T cell conjugates. Mucosal Immunol 3, 506–522 (2010).

19. Hladik, F., et al. Initial events in establishing vaginal entry and infection by human immunodeficiency virus type-1. Immunity 26, 257–270 (2007).

20. Patterson, B.K., et al. Susceptibility to human immunodeficiency virus-1 infection of human foreskin and cervical tissue grown in explant culture. Am J Pathol 161, 867–873 (2002).

21. Bertram, K.M., et al. Identification of HIV transmitting CD11c(+) human epidermal dendritic cells. Nat Commun 10, 2759 (2019).

22. Pena-Cruz, V., et al. HIV-1 replicates and persists in vaginal epithelial dendritic cells. J Clin Invest 128, 3439–3444 (2018).

23. Harman, A.N., et al. HIV-1-infected dendritic cells show 2 phases of gene expression changes, with lysosomal enzyme activity decreased during the second phase. Blood 114, 85–94 (2009).

24. Turville, S.G., et al. Immunodeficiency virus uptake, turnover, and 2-phase transfer in human dendritic cells. Blood 103, 2170–2179 (2004).

25. Haniffa, M., Collin, M. & Ginhoux, F. Identification of human tissue cross-presenting dendritic cells: A new target for cancer vaccines. Oncoimmunology 2, e23140 (2013).

26. Haniffa, M., et al. Human Tissues Contain CD141hi Cross-Presenting Dendritic Cells with Functional Homology to Mouse CD103+ Nonlymphoid Dendritic Cells. Immunity 37, 60–73 (2012).

27. Haniffa, M., et al. Differential rates of replacement of human dermal dendritic cells and macrophages during hematopoietic stem cell transplantation. J Exp Med 206, 371–385 (2009).

28. Turville, S.G., et al. Diversity of receptors binding HIV on dendritic cell subsets. Nat Immunol 3, 975–983 (2002).

29. Dominguez-Soto, A., et al. The DC-SIGN-related lectin LSECtin mediates antigen capture and pathogen binding by human myeloid cells. Blood 109, 5337–5345 (2007).

30. Dominguez-Soto, A., et al. Dendritic cell-specific ICAM-3-grabbing nonintegrin expression on M2-polarized and tumor-associated macrophages is macrophage-CSF dependent and enhanced by tumor-derived IL-6 and IL-10. J Immunol 186, 2192–2200 (2011).

31. Tang-Huau, T.L., et al. Human in vivo-generated monocyte-derived dendritic cells and macrophages cross-present antigens through a vacuolar pathway. Nat Commun 9, 2570 (2018).

32. Tang-Huau, T.L. & Segura, E. Human in vivo-differentiated monocyte-derived dendritic cells. Semin Cell Dev Biol 86, 44–49 (2019).

33. Cavarelli, M., Foglieni, C., Rescigno, M. & Scarlatti, G. R5 HIV-1 envelope attracts dendritic cells to cross the human intestinal epithelium and sample luminal virions via engagement of the CCR5. EMBO Mol Med 5, 776–794 (2013).

34. Gurney, K.B., et al. Binding and transfer of human immunodeficiency virus by DC-SIGN+ cells in human rectal mucosa. J Virol 79, 5762–5773 (2005).

35. Shen, R., Smythies, L.E., Clements, R.H., Novak, L. & Smith, P.D. Dendritic cells transmit HIV-1 through human small intestinal mucosa. J Leukoc Biol 87, 663–670 (2010).

36. McGovern, N., et al. Human Dermal CD14+ Cells Are a Transient Population of Monocyte-Derived Macrophages. Immunity 41, 465–477 (2014).

37. Goudot, C., et al. Aryl Hydrocarbon Receptor Controls Monocyte Differentiation into Dendritic Cells versus Macrophages. Immunity 47, 582–596 e586 (2017).

38. Segura, E., et al. Human inflammatory dendritic cells induce Th17 cell differentiation. Immunity 38, 336–348 (2013).

39. Patel, V.I., et al. Transcriptional Classification and Functional Characterization of Human Airway Macrophage and Dendritic Cell Subsets. J Immunol 198, 1183–1201 (2017).

40. Botting, R.A., et al. Phenotypic and functional consequences of different isolation protocols on skin mononuclear phagocytes. J Leukoc Biol 101, 1393–1403 (2017).

41. Harman, A.N., et al. HIV infection of dendritic cells subverts the interferon induction pathway via IRF1 and inhibits type 1 interferon production. Blood (2011).

42. Harman, A.N., et al. HIV Blocks Interferon Induction in Human Dendritic Cells and Macrophages by Dysregulation of TBK1. J Virol 89, 6575–6584 (2015).

43. Harman, A.N., et al. HIV induces maturation of monocyte-derived dendritic cells and Langerhans cells. J Immunol 177, 7103–7113 (2006).

44. Nasr, N., et al. HIV-1 infection of human macrophages directly induces viperin which inhibits viral production. Blood 120, 778–788 (2012).

45. Picelli, S., et al. Smart-seq2 for sensitive full-length transcriptome profiling in single cells. Nat Methods 10, 1096–1098 (2013).

46. Picelli, S., et al. Full-length RNA-seq from single cells using Smart-seq2. Nat Protoc 9, 171–181 (2014).

47. Kim, D., et al. TopHat2: accurate alignment of transcriptomes in the presence of insertions, deletions and gene fusions. Genome Biol 14, R36 (2013).

48. Xu, G., et al. SAMMate: a GUI tool for processing short read alignments in SAM/BAM format. Source Code Biol Med 6, 2 (2011).

49. McGovern, N., et al. Human dermal CD14(+) cells are a transient population of monocyte-derived macrophages. Immunity 41, 465–477 (2014).

50. Bigley, V., et al. Langerin-expressing dendritic cells in human tissues are related to CD1c+ dendritic cells and distinct from Langerhans cells and CD141high XCR1+ dendritic cells. Journal of Leukocyte Biology 97, 627–634 (2015).

51. Colomer-Lluch, M., Ruiz, A., Moris, A. & Prado, J.G. Restriction Factors: From Intrinsic Viral Restriction to Shaping Cellular Immunity Against HIV-1. Front Immunol 9, 2876 (2018).

52. Harman, A.N., et al. Identification of lineage relationships and novel markers of blood and skin human dendritic cells. J Immunol 190, 66–79 (2013).

53. Perot, B.P., Garcia-Paredes, V., Luka, M. & Menager, M.M. Dendritic Cell Maturation Regulates TSPAN7 Function in HIV-1 Transfer to CD4(+) T Lymphocytes. Front Cell Infect Microbiol 10, 70 (2020).

54. Ruffin, N., et al. Constitutive Siglec-1 expression confers susceptibility to HIV-1 infection of human dendritic cell precursors. Proc Natl Acad Sci U S A 116, 21685–21693 (2019).

55. Deymier, M.J., et al. Heterosexual Transmission of Subtype C HIV-1 Selects Consensus-Like Variants without Increased Replicative Capacity or Interferon-alpha Resistance. PLoS Pathog 11, e1005154 (2015).

56. Gringhuis, S.I., et al. HIV-1 exploits innate signaling by TLR8 and DC-SIGN for productive infection of dendritic cells. Nat Immunol 11, 419–426 (2010).

57. Garcia-Vallejo, J.J., et al. Glycodendrimers prevent HIV transmission via DC-SIGN on dendritic cells. Int Immunol 25, 221–233 (2013).

58. Cunningham, A.L., Harman, A.N. & Donaghy, H. DC-SIGN ‘AIDS’ HIV immune evasion and infection. Nat Immunol 8, 556–558 (2007).

59. Sewald, X., et al. Retroviruses use CD169-mediated trans-infection of permissive lymphocytes to establish infection. Science 350, 563–567 (2015).

60. Hammonds, J.E., et al. Siglec-1 initiates formation of the virus-containing compartment and enhances macrophage-to-T cell transmission of HIV-1. PLoS Pathog 13, e1006181 (2017).

61. Kashem, S.W., Haniffa, M. & Kaplan, D.H. Antigen-Presenting Cells in the Skin. Annu Rev Immunol 35, 469–499 (2017).

62. Stax, M.J., et al. Mucin 6 in seminal plasma binds DC-SIGN and potently blocks dendritic cell mediated transfer of HIV-1 to CD4(+) T-lymphocytes. Virology 391, 203–211 (2009).

63. Gringhuis, S.I., den Dunnen, J., Litjens, M., van der Vlist, M. & Geijtenbeek, T.B. Carbohydrate-specific signaling through the DC-SIGN signalosome tailors immunity to Mycobacterium tuberculosis, HIV-1 and Helicobacter pylori. Nat Immunol 10, 1081–1088 (2009).

64. van der Vlist, M., van der Aar, A.M., Gringhuis, S.I. & Geijtenbeek, T.B. Innate signaling in HIV-1 infection of dendritic cells. Curr Opin HIV AIDS 6, 348–352 (2011).

65. Guilliams, M., et al. Unsupervised High-Dimensional Analysis Aligns Dendritic Cells across Tissues and Species. Immunity 45, 669–684 (2016).

66. Burton, D.R., et al. Limited or no protection by weakly or nonneutralizing antibodies against vaginal SHIV challenge of macaques compared with a strongly neutralizing antibody. Proc Natl Acad Sci U S A 108, 11181–11186 (2011).

67. Botting, R.A., et al. Langerhans cells and sexual transmission of HIV and HSV. Rev Med Virol 27(2017).

68. van den Berg, L.M., et al. Caveolin-1 mediated uptake via langerin restricts HIV-1 infection in human Langerhans cells. Retrovirology 11, 123 (2014).

69. de Witte, L., et al. Langerin is a natural barrier to HIV-1 transmission by Langerhans cells. Nat Med 13, 367–371 (2007).

